# Combinatorial library of biodegradable polyesters enables delivery of plasmid DNA to polarized human RPE monolayers for retinal gene therapy

**DOI:** 10.1101/264390

**Authors:** Bibhudatta Mishra, David R. Wilson, Srinivas R. Sripathi, Mark P. Suprenant, Yuan Rui, Karl J. Wahlin, Cindy Berlinicke, Jordan J. Green, Donald J. Zack

## Abstract

Efficient gene delivery into hard-to-transfect cells is still a challenge despite significant progress in the development of various gene delivery tools. Non-viral and synthetic polymeric nanoparticles offer an array of advantages for gene delivery over the viral vectors and high in demand as they are safe to use, easy to synthesize and highly cell-type specific. Here we demonstrate the use of a high-throughput screening (HTS) platform to screen for biodegradable polymeric nanoparticles (NPs) that can transfect human retinal pigment epithelial (RPE) cells with high efficiency and low toxicity. These NPs can deliver plasmid DNA (pDNA) to RPE monolayers more efficiently compared to the commercially available transfection reagents without interfering the global gene expression profile of RPE cells. In this work, we have established an HTS platform and identified synthetic polymers that can be used for high efficacy non-viral gene delivery to human RPE monolayers, enabling gene loss- and gain-of-function studies of cell signaling and developmental pathways. This platform can be used to identify the optimum polymer, weight-to-weight ratio of polymer to DNA, and the dose of NP for various retinal cell types.

## INTRODUCTION

Gene therapy holds potential promise for treating both acquired and inherited blinding disorders as most of the identified disease gene to date is associated with RPE^1^. Modulating specific gene targets simply by turning off or turning on its function has become a standard tool to enhance stem cell differentiation or to reprogram induced pluripotent stem cells (iPSCs) from somatic cells ^2,3^. Routinely approached gene therapy utilizes viral vectors to deliver pDNA considering their potential for high-efficiency gene delivery. However, to its flip side, this approach is limited by several different factors such as (a) potential for insertional mutagenesis^4^, (b) prone to degradation in the cytosol by nucleases^5^ or can accommodate only a specific size of pDNA to deliver^6-8^. To overcome these challenges and to follow an alternative safer approach, significant attempts have been made to formulate and develop biodegradable non-viral vehicles agents to facilitate delivery of the gene of interest to the target sites.

As the charge distribution on both the plasmid DNA and the cellular membrane is profoundly negative, cationic polymers often demonstrate efficient intracellular delivery by merely condensing the cargo (pDNA) by strong electrostatic interaction and form NPs^9^. Since this strategy is episomal, it is usually considered as the safest way of delivering genes into the subcellular targets^10^. To this end, a range of different cationic polymers have been formulated and studied over the years for efficient non-viral gene delivery strategies^11-17^. Regardless of the advantages that cationic polymers demonstrate over the viral mode of gene delivery, it’s application is limited by a significant factor, i.e., poor transfection efficacy^18^. Poly(b-amino ester)s (PBAEs), a class of synthetic, cationic polymers, recently found to be useful as non-viral gene delivery agents. PBAEs are preferred polymers as they are easy to synthesize and demonstrate an efficient binding with its DNA counterpart. PBAEs are also hydrolytically degradable under physiological conditions and hence exhibit minimal cytotoxicity upon cellular administration. PBAEs have been shown in our prior work to be successful in transfecting human adult and embryonic stem cells^19^, and mouse RPE cells *in vitro* and *in vivo*^20^. Besides, prior works from our laboratory have also suggested PBAEs to have cell-type specificity based on their chemical structures^21,22^. Hence PBAEs makes an ideal carrier to undertake this study given their structural tenability and simple synthesis scheme.

The RPE cells are composed of a monolayer of pigmented and bipolar epithelial cells at the back side of the retina. Any compromise in the cellular environment of RPE cells leads to many hereditary and acquired diseases, including age-related macular degeneration (AMD)^23^. As RPE also dispensable for photoreceptor turnover and maintenance and as both PR and RPE dominate the retinal cell population, RPE cells could be the targets of therapy in many ocular diseases. Moreover, as in many ocular diseases lead to overall genetic imbalance^24,25^, gene therapy is vital in restoring the gene expression in the compromised retina. Attempt to deliver gene either into primary RPE cells or RPE cell lines is not new in the field. However, despite adopting several different non-viral strategies to deliver DNA either by polymeric or by liposomal vectors, the success rate is very low^26-35^. In this study, we have established a high throughput-screening platform to screen for potential PBAE nanoparticles to access its transfection efficacy in iPS derived human RPE cells *in vitro.* We hypothesize that cationic PBAE- pDNA NP complex can be delivered to the RPE monolayer efficiency by tuning the hydrophilicity and end group chemistry. To test our hypothesis, we synthesized a library of 4 PBAE base polymers with different backbone and end-group chemistry. The ability of PBAE to bind to its DNA counterpart was examined by electrophoresis assay. To explore the effects of different PBAE chemical structures on delivery efficiency, we transfected 25 days old RPE monolayers with 140 different combinations of PBAEs using a pDNA encoding mCherry reporter gene under the CAG promoter. The outcomes were evaluated in a High Content Analysis platform where the images were acquired, and the data analysis was performed using specific algorithms.

## RESULT

### Polymer Synthesis

In the very first step, to achieve successful transfection, we intended to formulate a group of stable nanoparticles. NP formulation with the different combination of end-capped polymer and pDNA occurs via strong electrostatic interaction. In this current work, we have used two different plasmids expressing either the mCherry reporter or the nuc-GFP reporter driven by the same CMV early enhancer/chicken β actin (CAG) heterologous promoter as described in material and method section for the HTS. Once the linear PBAE synthesis was completed, all the subsequent steps including end-capping reaction, preparation of source plate with end-capped polymers, stable NP formation with the desired pDNA, automated dispensing for transfection, and the HCA image capturing processes were carried out in a 384 well format for all the combination of NPs **(Schematic-1)**. A range of different base polymers end-capped with a variety of different amino-terminal structures was combined to prepare a combinatorial library of 144 different PBAE NP formulations. The polymer nomenclature “N1-N2-XN” in the whole library, denotes base polymer number (N)-side chain number (N)- end cap amino terminal type (X) and number (N) respectively **(Figure-1)**.

**Figure 1:**
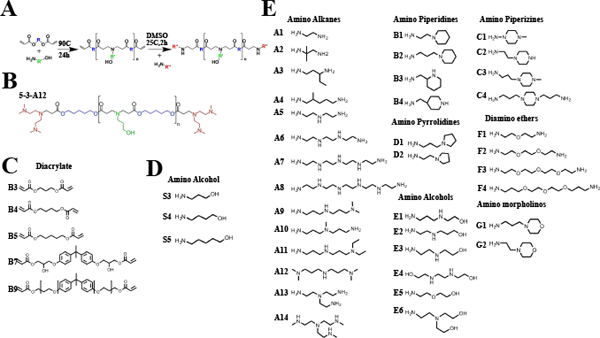
Sequential poly(beta-amino ester)s (PBAEs) library construction and synthesis scheme. **(A)** Synthesis scheme of linear PBAEs from diacrylate and primary amine small monomers to yield acrylate terminated polymers followed by end-capping to yield linear end-capped PBAEs. **(B)** Example PBAE 5-3-A12 formed from monomers B5, S3 and end-cap A12. **(C)** Five diacrylate monomers and **(D)** three side-chain amino alcohols utilized in library synthesis. **(E)** 36 end-cap monomers identified as effective for transfection.

### High Throughput automated NP transfection to RPE monolayer

To access the transfection efficacy of the PBAE/pCAGG-mCherry nanoparticles in matured RPE monolayers (Day 25 post seeding), we conducted a high throughput-screening assay with all 144 different combinations of nanoparticles as explained in schematic-1. This allowed us a direct visualization of transfection efficacy **(Figure 2A)** and viability rate **(Figure 2B)** on a HCA platform where we collected images and analyzed data using the in built specific algorithm suitable to measure either transfection efficacy or viability rate. Cells transfected with the polymer without any end-capping reaction were included as controls. Heat map suggests that transfection efficacy differs significantly depending upon the side-chain end-capping chemistry of the PBAE **(Figure 2A)**. A few leading PBAE structures 5-3-A12, 5-3-F3 and 5-3-F4 resulted in 42%, 37% and, 34% positively transfected cells respectively. Interestingly, these specific polymers also demonstrated a significantly higher cell survival rate (90%, 97% and, 98% respectively; **figure 2B**). However, cell survival property of these top polymers was not directly proportional to their ability to transfect RPE monolayers, as some other PBAEs demonstrated extremely low transfection efficacy irrespective of their high cell survival property. We also noticed that different PBAEs pairs with same end-capping molecules shown substantially different transfection efficiency, suggesting that the transfection efficiency is also dependent upon additional parameters such as the degree of hydrophilicity and overall NP stability (e.g., the transfection efficacy of 3-5-A12 is 3.8% while it is 42% for 5-3-A12). In addition, we also noticed that, while the PBAE 5-3-A12 demonstrated highest transfection efficiency (42%) and higher survival rate (90%) to the monolayered RPE cells at day 25, the same polymer yielded a lower transfection rate (33%) and lower survival rate (30%) in differentiating RPE cells at the early phase of differentiation on day 3 **(Figure S2)**. This suggests that the overall transfection efficacy and impact on cell survival rate of a particular formulation varies significantly between different phases of a “differentiating” RPE cell.

**Figure 2:**
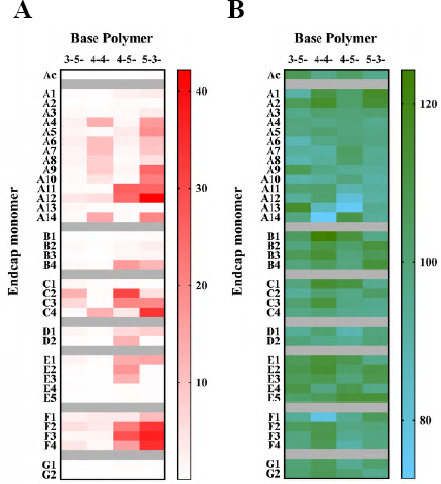
In vitro high throughput screening of PBAE nanoparticles in confluent D25 RPE monolayer. **(A)** Heat maps showing the percentage transfected RPE cells and **(B)** percentage survival rate following the introduction of a combinations of 140 different nanoparticles to confluent RPE monolayer at day 25 post seeding. The color scale bar refers to the percentage transfection efficiency and percentage survival that was calculated based on the number of mCherry positive cells detected from total number of cell population.

### Biophysical characterization of 5-3-A12 nanoparticle

To further investigate the biophysical properties of PBAE nanoparticles that demonstrated high efficiency of pCAGG-mCherry delivery, we measured the particle size of the 5-3-A12 nanoparticle by both dynamic light scattering (DLS) and nanoparticle tracking analysis (NTA) methods and also measured zeta potential. All parameters were measured at different weight-to-weight (w/w) ratio. The particle size demonstrated a fairly broad distribution ranging from 49 nm to 191 nm by DLS method, and from 115 nm to 149 nm by NTA method **(Figure 3A, 3B)**. In corroboration with a previous report our results suggest that nanoparticle with the smaller size results in increased transfection rate^36^, as during the transfection optimization process, we always noticed higher transfection rate at lower w/w ratio compared to when we used higher w/w ratio. Regardless, the transfection efficiency was always higher with the 5-3-A12 nanoparticle at any w/w ratio compared to other lead nanoparticles. Given the comparable transfection efficiency of the 5-3-A12 nanoparticle with different particle size, our results suggest that transfection efficiency of the 5-3-A12 nanoparticle is not solely dominated by particle size. Different than the particle size, 5-3-A12 nanoparticle demonstrated fairly similar surface charge distribution at any given size, which ranges from +25 mV to +30 mV as measured by zeta potential **(Figure 3C)**. Gel electrophoresis study demonstrated a complete PBAE/ pCAGG-mCherry nanoparticle complex formation **(Figure 3D)**. To further consolidate the biophysical results observed, we conducted a transmission electron microscope (TEM) analysis. TEM imaging confirmed stable PBAE/ pCAGG- mCherry nanoparticle formation through the self-assembly process, with nanoparticle size consistent with the DLS and NTA results **(Figure 3E)**.

**Figure 3:**
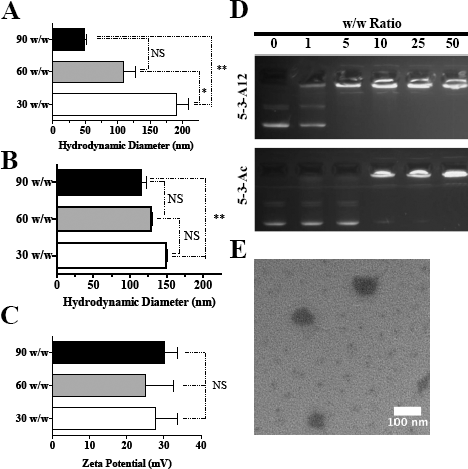
PBAE 5-3-A12 Characterization. PBAE 5-3-A12 characterization. **(A)** Diameter measurements assessed via DLS z-average and **(B)** NTA showed that average diameter decreased as polymer: DNA w/w ratio increased. DLS z-average measurements were statistically lower for 90 w/w nanoparticles, compared to 30 w/w nanoparticles **(C)** Nanoparticle zeta-potential did not statistically differ between the nanoparticles at different w/w ratios. **(D)** End-capping with monomer A12 improved DNA binding compared to acrylate-terminated polymers. PBAE 5-3-A12 fully retarded DNA at w/w ratios down to 5 w/w, in contrast to the acrylate terminated polymer, which was only effective down to a 10 w/w ratio. **(E)** TEM showed 5-3-A12 nanoparticles as spherical Graphs show mean of three independently prepared samples. *p < 0.01, *\*\*p* < 0.001, based on one-way ANOVA with Tukey's*post hoc* test.

### Validation of 5-3-A12 nanoparticle transfection efficacy

To further examine and validate the ability of the lead PBAE/pCAGG-mCherry nanoparticles to transfect RPE monolayers, 25 days old RPE monolayers were transfected separately in a 8-well chambered cover glass using already optimized transfection condition. RPE monolayers were also transfected with Lipofectamine 3000 and DNA-In for to test (and compare) the transfection efficacy as a control. We observed transfection efficiency up to 42% with the lead nanoparticle (5-3-A12), which were 26% higher than that of DNA-In and 41% higher than Lipofectamine-3000 as well as comparable transfection efficiency of all the other polymers **(Figure 4A-B)**. Interestingly, although 5- 3-A12 achieved the highest transfection in human RPE monolayers, it was the most inefficient polymer for mouse photoreceptor and human Retinal Ganglion Cells (data not shown) suggesting for its cell type specificity and suitable only for human RPE monolayers. To ensures that the differences in transfection efficiency were not caused by potential toxicity from polymeric nanoparticles, we also examined the effects of PBAE/pCAGG-mCherry nanoparticles on cell viability under the optimized transfection doses. Our results showed most PBAE/pCAGG-mCherry nanoparticles formulations did not negatively affect cell viability compared to untreated cells alone **(Figure 4C)**. The only exception was one of the commercial reagents DNA-In, which showed slightly lower cell viability (~80%). Previous work from our lab has reported a higher dose of PBAE required for the plasmid DNA delivery with weight ratios up to 50:1 to reach optimal transfection efficiency in cancer cells, which may also cause increased cell death^22^. However, in this study, substantially less PBAE was required (3:1) to form stable nanoparticles with top PBAEs due to the smaller size. Also, because of the cell type specificity, we observed minimal cell death with RPE monolayers. This offers an additional advantage of using PBAE for pDNA delivery given the minimal toxicity effects with different cells. We also quantified mean fluorescent intensity, which is a measure of total protein production. In this regard, RPE monolayers transfected in any form (via PBAEs or via commercial reagents) demonstrated a substantially similar level of mCherry intensity, despite their comparable level of percentage of cells being transfected **(Figure 4D)**. Transfection efficiency describes the percentage of cells that have been transfected, regardless of the difference in the level of protein production among individual cells. In contrast, mean fluorescence intensity takes into account of the difference in protein production by individual cells, and normalize that by the total number of cells. Therefore, mean fluorescence intensity is a better prediction of the level of protein production post-transfection. Also, we also counted the total number of cells over time during the differentiation process and measured relative cell count and transfection efficacy of lipofectamine and DNA-In on RPE monolayers at different DNA doses. Even though the cell number over time and the overall post-transfection viability rate was acceptable, the transfection efficacy was weak compared to 5-3-A12 PBAE at any given DNA dose **(Figure S4)**.

**Figure 4:**
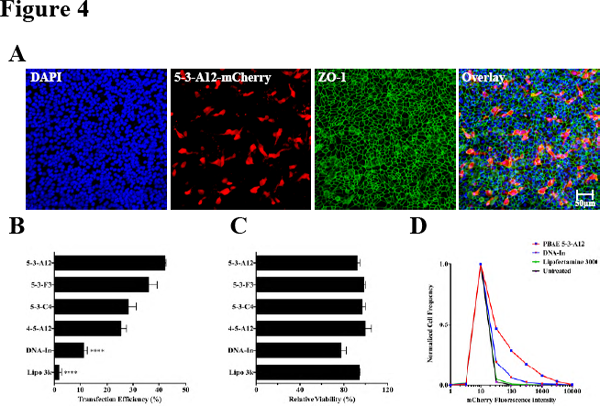
*in vitro* transfection of confluent D25 RPE monolayer with top PBAE nanoparticles hits obtained from the preliminary high throughput screening. **(A)**Representative Z-stack confocal micrographs showing transfected RPE cells (red) and the epical localization of ZO-1 protein (green) of RPE monolayers transfected with a PBAE nanoparticle (5-3-A12) that yielded highest transfection efficacy in D25 RPE monolayer. The nuclei were counterstained with DAPI (blue). Histogram showing **(B)** transfection efficacy **(C)** relative viability and **(D)** mean fluorescent intensity of top 3 hits obtained from preliminary screen (5-3-A12, 5-3-F3 and 5-3-F4) along with commercial transfection reagents (lipofectamine 3000 and DNA-In). Transfection efficiency shown by percentage of mCherry positive cells, quantified using a specific algorithm designed for transfection assay in a High Content Analysis platform. *\*\*\*\*p* < 0.001, based on student t-test. Confocal micrograph Scale bar: 50μm.

### Multiple gene delivery with 5-3-A12 nanoparticle into RPE monolayers

Since multiple gene deliveries using nanoparticle is quite challenging and none of the previous studies report using PBAE nanoparticle for multiple gene deliveries, we test the transfection efficacy of 5-3-A12 polymers for the delivery of more than one gene into RPE monolayers. To this end, we used two separate pDNA constructs encoding two different reporter genes (mCherry and nuclear GFP) under same promoter (CAGG) and optimized a co-transfection assay. We generated comparative data for cells that received either one or both of the reporter genes in a co-transfected cell population, 48 hours posttransfection **(Figure 5A)**. We adopted two different strategies for transfection; either both the constructs were transfected at the same time (Co-transfected) or at the different times (serially transfected). Post-transfection data was analyzed for transfection efficacy **(Figure 5B)**, cell body area **(Figure 5C)**, and cell body shape **(Figure 5D)** for each condition. Our data suggest that under co-transfection condition, about 50% of the cell population received NP containing mCherry pDNA and about 25% of the cell population received NP containing nuc-GFP pDNA and remaining 25% cells obtained both the plasmids. In contrary, serially transfected condition favored more to NP containing mCherry pDNA, where more than 97% cell population received NP containing mCherry pDNA. While the preference of receiving one plasmid over another was significantly different in both the transfection conditions, as expected, we noticed no apparent noticeable change either in cell body shape or cell body size in either circumstances. These results suggest that 5-3-A12 polymers do not interfere with intrinsic cellular pathways that trigger cellular/nuclear morphology and encourages for non-viral gene delivery applications.

**Figure 5:**
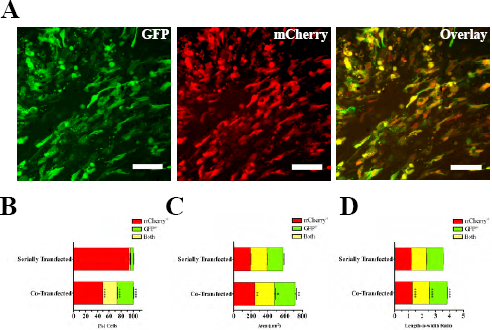
Transfection efficacy as measured in a co-transfection assay. **(A)** Representative Cellomics images of RPE monolayers co-transfected with both mCherry (red) and GFP (green) constructs. Histogram showing **(B)** *%* cells **(C)** cell body area and **(D)** cell body size of cells that were introduced with either mCherry or GFP alone or cotransfected with both the construct.

### Effect of 5-3-A12 NP transfection on RPE markers gene expression

Next, we directly measured endogenous RPE known marker gene expression from 5-3- A12 NP transfected population. In this assay, we wanted to evaluate if the 5-3-A12 NP transfection is altering any RPE marker gene expression. RNA samples were harvested, 48 hours post-transfection and flow sorted to collect mCherry+ and mCherry_-_ cell population. RNA samples were also harvested from the un-transfected cells as a positive control **(Figure 6A)**. RPE specific gene expression changes were then compared between different samples obtained from qPCR experiment **(Figure 6B)**. To our surprise, we found subtle alteration in gene expression from the samples collected from transfected wells (irrespective of transfection) compared to the samples from untransfected wells. While the fold change of expression level of *Mitf, Ncad* and, *Otx2* were increased in both mCherry+ and mCherry^-^ cell population, the expression level of *Sox9, Tyr, Rpe65, R1bp1, Pmel17* and *Best 1* were decreased in both the population. While *Lrat* expression was decreased in mCherry+ population, it was 0.5 fold higher in mCherry^-^ cell population. In contrary, while *Crx* expression was decreased in mCherry^-^ population, it was slightly higher (0.3 fold) in mCherry+ cell population. *Ecad* expression was slightly decreased in mCherry+ population, and it was similar to control level in mCherry^-^ cell population. Even though all the changes we noticed are negligible, we speculate that the changes are transient with respect to the stress level that the cells receive from the transfection procedure.

**Figure 6:**
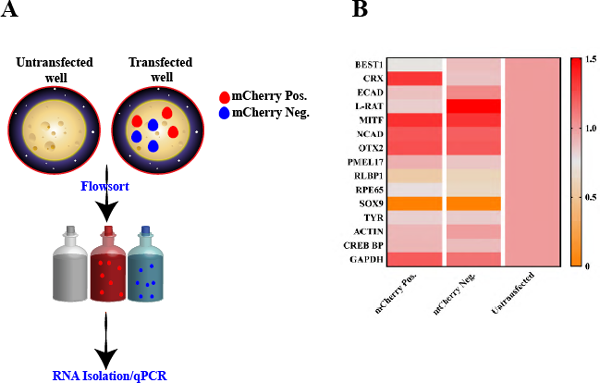
Post transfection RPE Specific gene expression profile. **(A)** schematic showing a flow cytometry assay to collect mCherry positive and mCherry negative cells from a transfected cell population **(B)** RPE specific gene expression in flowsorted hiPSC- RPE as quantified by qPCR assay. Error bars represent SD of biological replicates. BEST1 (bestrophin 1), PMEL17 (premelanosome protein), RPE65 (retinal pigment epithelium-specific protein), TYR (tyrosinase), CRX (Cone-Rod Homeobox protein), E- CAD (E-Cadherin), L-RAT (lecithin retinol acetyltransferase), MITF (melanogenesis associated transcription factor), N-CAD (E-Cadherin), OTX2 (orthodenticle homeobox- 2), RLBP-1 (retinaldehyde-binding protein 1), SOX-9 (SRY-Box9), Actin, CREB BP (cAMP response element binding protein), GAPDH (Glyceraldehyde 3-phosphate dehydrogenase).

## DISCUSSION

The most successful *in vitro* plasmid DNA gene delivery studies were established on RPE-derived cell lines, which are easier to transfect than primary RPE monolayers37. In this study, we have used hiPSc-derived RPE cells, as they are considered to be much more similar to primary RPE than RPE cell lines^38^ to investigate the utility of using a biodegradable and non-viral gene delivery approach for transient protein expression in primary RPE monolayers. To this end, we have established a high-throughput platform to screen NPs created from a wide variety of polymers for their ability to deliver gene into the human stem cell-derived RPE monolayers. Using this system, we have identified synthetic polymers that can be used for high efficacy non-viral gene delivery to human RPE monolayers, enabling gene loss- and gain-of-function studies of cell signaling and developmental pathways. Since the self assemble process of polymers high very complex^39^, it is very important to combine appropriate physical, chemical and biological properties to produce efficient polymers for gene delivery. Thus, high-throughput parallel generation and screening of large libraries of such nano-carriers is a very efficient and powerful way to identify efficacious and non-toxic gene delivery vectors.

Despite the great interest in hiPSC RPE cells as sources for cell therapy and *in vitro* disease modeling, no studies of gene delivery of these cells using PBAE nanoparticles have, to our knowledge, been reported. We have shown that, as is the case for RPE cells, hiPSC -RPE cells are very difficult to transfect with plasmid DNA complexed with any commercial transfection reagent (lipofectamine or DNA-In). The highest efficiency of transfection with plasmid DNA using DNA-In was achieved on RPE monolayers with an efficiency of about 10%, and this was even lower (less than 5%) with lipofectamine 3000. In addition, in our high-throughput screening assay, while most of the PBAE demonstrated a decent range of transfection (~10% to ~50%) on the sub-confluent RPE population (day 3 after seeding), most of them fail to transfect confluent RPE monolayer population (day 25 after seeding) when the cells reached a polygonal morphology. In contrast, top hits from the screening (5-3-A12, 5-3-F3 and 5-3-F4) could able to deliver the pDNA efficiently into both sub-confluent, and post-confluent monolayer (polygonal) RPE cells. While the reason for this discrepancy is not clear, we speculate that it is a phase dependent cell-type-specific event where the preferences of the interaction of the cationic polymer changes with the cell membrane structural change over time. Further optimization studies with different transfection agents, ratios of modified PBAEs, and pDNA/PBAE ratios are needed to understand this. Regardless, our results suggest that 5- 3-A12 PBAE nanoparticle met all the criteria of a successful non-viral gene therapy agent being readily internalized into the cell, escaped endocytic degradation and successfully delivered the pDNA into the nucleus to be expressed. Even though we did not undertake any specific characterization of the uptake of PBAE NPs *in vitro;* however our confocal imaging data with ZO-1 labeling indicate that exclusively RPE monolayers with polygonal shape take up the particles. In our co-transfection assay, while the preference for getting transfected either with pmCherry or pNucGFP was subtle, our result from serially transfected RPE monolayer suggests that already transfected cells are less receptive or more rigid for re-uptake of new nanoparticles. This conclusion is based on the fact that in serially transfected cells the mCherry transfected cell population outnumbered dramatically to that of GFP transfected cell population when the cells were transfected with mCherry construct first. However, regardless of the type of transfection, PBAE nanoparticle has no impact on either cell body shape or size as evident from our co-transfection assay. This observation also suggests that, although PBAE nanoparticles can deliver multiple genes into RPE monolayers, they are often hampered by poor reproducibility and low co-transfection efficiency especially when cells are transfected serially. Our results also suggest that the PBAE nanoparticle 5-3-A12 can preferentially deliver pDNA into human RPE monolayers with relatively low cytotoxicity. Even though the mechanism-of-action (MoA) is not known at this time, results from the current work provides important insights and holds promises for translational application of the biodegradable PBAE nanoparticles especially for RPE dysfunction. Since the overall surface charge distribution is an important deciding factor on cellular cytotoxicity^40,41^, we speculate two different theories that results in low cytotoxicity effect of 5-3-A12 nanoparticle. (1) its overall charge distribution on the surface (ranges from +25 mV to +30 mV) at any given w/w/ ratio, which helps in interacting with the negatively charged components at the cell surface and destabilizes the cell membrane more efficiently than any other polymers used during primary screening and (2) the electrostatic interaction between 5-3-A12 and pDNA introduces sufficient number of available amine group in 5- 3-A12 that could results in increased zeta potential value.

In this work we have validated the expression pattern of known RPE markers from from both mCherry^+^ and mCherry^−^ cell population by low throughput (96-well) format using a qRT PCR assay. This purpose was to evaluate the possible PBAE interference with any known intrinsic RPE gene pathway. We were not expecting any change in the gene expression pattern after transfection as the pDNA used expresses exogenous reporter genes without any known function on RPE markers. However, we found differential gene expression pattern from the sample collected from transfected wells (regardless of their transfection status) compared to sample collected from the un-transfected wells.

## MATERIALS AND METHODS

### Polymer synthesis and characterization

Monomers were purchased from vendors listed in Supplemental Table 1 and Supplemental Table 2. Acrylate monomers were stored with desiccant at 4°C, while amine monomers were stored with desiccant at room temperature. PBAE polymers were synthesized neat at 1.1:1 B:S monomer ratios for polymers 3-5-Ac, 4-4-Ac and 4-5-Ac and 1:1.05 monomer ratio for 5-3-Ac for 24 hours at 90°C. Following synthesis, neat polymers were dissolved at 200 mg/mL in anhydrous DMSO then precipitated in diethyl ether twice at a solvent ratio of 1:10 by vortexing the solvents and centrifuging at 3000 rcf. Polymers were allowed to dry under vacuum for 24 hours, at which point they were massed and dissolved at 200 mg/mL in anhydrous DMSO and allowed to remain under vacuum to remove additional diethyl ether for another 24 hours. Finally, acrylate terminated polymers were aliquoted and stored at -20°C until use in end capping reactions.

For polymer characterization, samples of the initial neat polymer and neat polymer following diethyl ether removal were set aside for characterization via ^1^H NMR and gel permeation chromatography (GPC). GPC was performed on polymer samples both before and after double precipitation in diethyl ether using a Waters system with auto sampler, styragel column and refractive index detector to determine MN, MW and PDI relative to linear polystyrene standards. GPC measurements were performed as previously described with minor changes of a flow rate (0.5 mL/min) and increase in sample run time to 75 minutes per sample^42^. Analysis of polymers via ^1^H NMR (Bruker 500 MHz) following diethyl ether precipitation and drying was performed to confirm the presence of acrylate peaks. For NMR, neat polymer was dissolved in CDCl_3_ containing 0.05% v/v tetramethylsilane (TMS) as an internal standard.

### Polymer library preparation

PBAE polymers were prepared for transfection screening experiments by high-throughput, semi-automated synthesis techniques using ViaFlo 384 (Schematic 1B). For end capping reactions, 25 μL of endcap molecules in anhydrous DMSO at a concentration of 0.2 M were distributed to source wells of a deep-well 384 well plate, then distributed to corresponding replicate wells in groups shown in multiple colors of the end capping reaction 384-well deep plate (240 μL volume). Acrylate terminated base polymers at 200 mg/μL in anhydrous DMSO were thawed and distributed to wells containing 36 different endcap molecules and a single well containing DMSO only for the acrylate terminated polymer control. End capping reactions were allowed to proceed for two hours at room temperature on a gentle shaker, after which endcapped PBAE polymers were diluted to 50 mg/mL in anhydrous DMSO and aliquoted to 5 μL per well on the left side of 384-well nanoparticle source plates. Nanoparticle source plates were sealed and stored at -20°C with desiccant until needed for transfection. Following large-scale screening of the PBAE library in 384 well plates, larger batches of top PBAE structures were synthesized from frozen base polymer using the same protocol described above. Endcapped polymers were then aliquoted to individual tubes and stored at -20°C with desiccant.

**Schematic 1:**
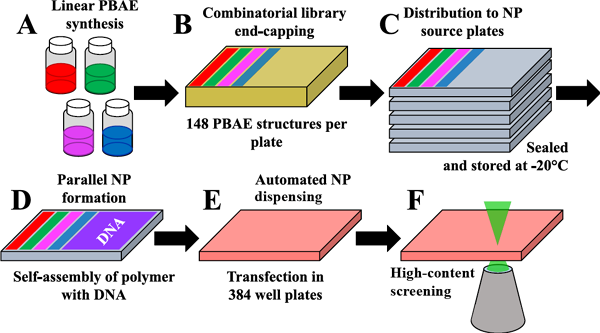
Schematic of combinatorial PBAE library construction. **(A)** Linear base polymer PBAEs were synthesized in vials to be acrylate terminated, then characterized via IH NMR and GPC **(B)** Synthesized polymers are dispensed into a 384 well round bottom plate using Viaflo 96/384 microplate dispenser and end-capped with each base polymer. A total of 4 different base polymers as shown in different color scheme are end capped per master plate containing 36 end-cap monomers each. **(C)** Source plates were then replicated from one master plate and stored them at -80C for future use. **(D)** End capped linear polymers (left 12 columns of the plate) were mixed with plasmid DNA (right 12 columns of the plate) to formulate NPs. **(E)** The RPE monolayers were transfected using automated Viaflo microplate dispenser and incubated for 48 hours with NPs. **(F)** Images were captured using Cellomics.

For end capping, reaction volumes of 50 μL at 100 mg/mL polymer concentration and 0.1 M were selected as sufficient to enable effective reactivity over a two-hour time period. For initial studies, endcap molecule E1 was titrated between 0.2 and 0.0625 M in reactions with base polymer PBAE 4-5-Ac at 100 mg/mL over two hours. Reacted polymers were then precipitated twice in diethyl ether to remove excess endcap monomer, dried and assessed using 1H NMR to determine efficacy of the end capping reaction by the disappearance of acrylate moiety peaks between 5.5-6.5 ppm. These results demonstrated effective end capping down to a concentration of 0.05 M for endcap molecule E1. To allow for varying levels of reactivity between endcap molecules, an endcap molecule concentration of 0.1 M was used for parallel large-scale end capping reactions.

### Nanoparticle characterization

The hydrodynamic diameter of top PBAE structure 5-3-A12 was characterized at three different w/w ratios to assess the influence of w/w ratio on nanoparticle characteristics. For dynamic light scatter (DLS) measurements, nanoparticles were initially formed in 25 mM NaAc, pH 5.0 then diluted 1:6 into 10% FBS in PBS dynamics and analyzed in disposable micro-cuvettes using a Malvern Zetasizer NanoZS (Malvern Instruments, Marlvern, UK) with a detection angle of 173°. For zeta potential, nanoparticles were prepared and diluted as for DLS, but were analyzed by electrophoretic light scattering was in disposable zeta cuvettes at 25°C using the same Malvern Zetasizer NanoZS. For nanoparticle tracking analysis, nanoparticles were formed in 25 mM NaAc, pH 5, then diluted 1:500 in 150 mM PBS as previously described using a Nanosight NS300.

A gel retention assay to assess PBAE: DNA binding strength was performed as previously described^43^ using a 1% agarose gel. Acrylate terminated PBAE 5-3-Ac was compared against top PBAE structure 5-3-A12 at w/w ratios from 0 to 50 to demonstrate improved binding of endcapped PBAE structures.

Transmission electron microscopy (TEM) images were acquired using a Philips CM120 (Philips Research, BriarcliffsManor, New York) on 400 square mesh carbon coated TEM grids. Samples were prepared at a DNA concentration of 0.045 μg/μL and polymer 90 w/w ratio in 25 mM NaAc, pH 5.0 after which 30 μL were allowed to coat TEM grids for 20 minutes. Grids were then dipped briefly in ultrapure water, wicked dry and allowed to fully dry before imaging.

### pDNA Design

For the *in vitro* transfection, a plasmid coding for the mCherry open reading frame was created by PCR amplification of the mCherry-N1 plasmid (Catalog no. 632523; Clontech). Since this plasmid has no start site an initiator, an ATG was added to the forward primer. After PCR amplification, mCherry was inserted into the directional pENTR-D-TOPO gateway entry vector (catalog no. K240020; Invitrogen). Positive colonies were selected by PCR and confirmed by sequencing. 100ng of purified entry plasmid was mixed with pCAGG-DV destination vector, created by incorporating a gateway cassette containing attR recombination sites flanking a ccdB gene into the pCAGEN vector (Addgene #11160), in the presence of LR clonase II (catalog no. 11791019). After recombination clones were selected and sequenced.

### Differentiation and Culture of RPE From hPSCs

RPE monolayers were differentiated as described previously by our laboratory^44,45^ from the EP1-GFP human iPS cell line that constitutively expresses H2B-nuclear-GFP. In brief, iPS cells to be differentiated were then plated at 60,000 cells per cm^2^ on Matrigel-coated 384 well plates and allowed to grow for 25 days in RPE medium consisting of 70% DMEM (catalog no. 11965092; ThermoFisher Scientific), 30% Ham’s F-12^46^ Nutrient Mix (catalog no. 11765-054; Invitrogen), serum free B27 supplement (catalog no. 17504044; ThermoFisher Scientific), and antibiotic-antimycotic (catalog no. 15240062; ThermoFisher Scientific). Coating of plate with Matrigel (25 μL per well), seeding of cells (50μL per well), and media change every other day (replaced with fresh 25μL per well) were accomplished using a high throughput Viaflo microplate dispenser (catalog no. 6031; Intergra). Cells were confirmed to possess an RPE monolayer phenotype at day 25 following plating.

### *In vitro* nanoparticle mediated gene delivery

On the day of transfection, the old media was discarded and replaced with 25μL of fresh RPE media. To form PBAE/DNA nanoparticles, pDNA was diluted in 25 mM sodium acetate buffer (NaAc, pH 5) and aliquoted to individual wells on the right half of the 384- nanoparticle-source plate. End capped PBAEs from the left half of the 384 well round bottom source well place (schematic figure-1D) were then resuspended in parallel in 25 mM NaAc using a Viaflo microplate dispenser. After a brief centrifugation (1000 rcf for 1 minute) the solutions of unique PBAE structures were then transferred to the right half of the 384 well round bottom source well place containing pDNA (schematic figure-1D) in a 3:1 (vol/vol) ratio, resulting in a defined weight-weight (w/w) ratio between 20-100 of PBAE:DNA. The nanoparticle source plate containing the PBAE/DNA mixtures was then briefly centrifuged (1000 rcf for 1 minute). To dispense nanoparticles to cells, 5 μL volumes of the NPs in each well were then added to the RPE monolayer (schematic figure-1E) and incubated with cells for 2 hours inside the 37°C incubator; all nanoparticles and media were then replaced with 50 μL of fresh RPE media. After 48 hours to allow for reporter gene expression, nuclei were stained with Hoechst and images acquired using an automated fluorescence-based imaging system (HCA Cellomics VTI; Thermofisher scientific). Transfected cells were identified as those expressing both the endogenous nuclear GFP and mCherry and the percent of transfected cells, as well as cell viability, was determined for each NP and condition.

Commercial transfection reagents Lipofectamine 3000^®^ (catalog no. L3000001; ThermoFisher Scientific) and DNA-In Stem (catalog no. GST-2130; MTI-Globalstem) were prepared according to manufacturer recommendations with pCAGG-mCherry. After particle formation, particles were added to day 25 differentiated RPE monolayer cells in 384 well plates at the specified DNA doses. Both reagents were optimized at multiple reagent:DNA ratios and for incubation times with cells of two hours and 24 hours to identify the optimal condition. After either two or 24 hours, media was entirely replaced with fresh medium and cells were cultured for two additional days, at which point transfection efficacy was assessed by image analysis with Cellomics.

### Immunostaining

iPS cells to be differentiated were plated at 2.3 million cells per cm^2^ on Matrigel-coated borosilicate sterile 8-well chambered cover glass (catalog no. 155409; Lab-Tek II;) and allowed to grow for 25 days in RPE medium. On the day of transfection, the old media was discarded and replaced with 300μL of fresh RPE media. The PBAE 5-3-A12 were then mixed with CAGG mCherry in a 3:1 (vol/vol) ratio, resulting in a defined weight-weight (w/w) ratio of 80:1 of PBAE:DNA. The nanoparticle containing the 5-3-A12 / CAGG mCherry mixtures was then briefly centrifuged (1000 rcf for 1 minute). To dispense nanoparticles to cells, 50 μL volumes of the NPs containing 1500 ng DNA were then added to the RPE monolayer and incubated with cells for 2 hours inside the 37°C incubator; all nanoparticles and media were then replaced with 300 μL of fresh RPE media. After 48 hours to allow for reporter gene expression the cells were fixed with 4% paraformaldehyde in PBS, cells were blocked and permeabilized for 30 min in 5% goat serum, 0.25% Triton X-100 in PBS, and then incubated for 1 h at room temperature with polyclonal mouse anti-ZO-1 (1/500; catalog no. 40-2200; Invitrogen) monoclonal rat anti-mCherry (1/1000; catalog no. M-11217; Molecular Probes). Cells were then incubated for 1 h at room temperature with the corresponding secondary antibody conjugated to Alexa 488 or Alexa 568 (Invitrogen), and counterstained with Hoechst 33342 (Invitrogen). Images were captured with a confocal microscope (Zeiss LSM 710).

### Co-expression Assay

To assess the ability of top PBAE nanoparticles to co-deliver two plasmids, EP1 cells that lacked nuclear GFP expression were plated as described above in 384 well plates and differentiated for 25 days to RPE monolayers. Plasmids CAGG-mCherry and CAGG- nucGFP were diluted in 25 mM NaAc as described above and used to form PBAE 5-3- A12 nanoparticles at an 80 w/w ratio and DNA dose of 200 ng/well in 384 well plates. For the co-delivered condition, plasmids in 25 mM NaAc were pre-mixed prior to complexation with PBAE and added to RPE monolayers together in the same nanoparticles. For the serial transfection experiment, nanoparticles formed with plasmid CAG-mCherry only were added to cells at a dose of 100 ng/well on day 25 following plating and nanoparticles containing plasmid CAG-GFP only were added to cells on day 26. Media changes were performed as described above. Transfection efficacy for GFP and mCherry was assessed on day 28 following staining of cell nuclei with Hoechst 33342.

### RT-qPCR Assay

Differentiated monolayer RPE cells in a 96 well plate were transfected as described above with PBAE 5-3-A12 nanoparticles at a surface area scaled dose. Following approximately 48 hours to allow for expression, cells were incubated with 0.25% trypsin and lifted from the plate surface to a single cell suspension. Trypsin was quenched with 2% FBS in PBS and cells were pelleted, resuspended in fresh 2% FBS buffer and sorted for mCherry fluorescence using a Sony SH800 flow sorter to positive and negative expressing populations. RNA was extracted from the positive and negative mCherry expressing cell populations using Qiagen RNAse Easy minikit (Catalog no. 74104; Qiagen) and reverse-transcribed (catalog number. 4368814; High Capacity cDNA kit; Applied Biosystem). Untreated wells were likewise trypsinized and processed prior to RNA extraction. Quantitative PCR samples were run in biological triplicates and expression levels normalized using the geometric mean of three reference genes: ACTIN, GAPDH, and CREBBP. Gene-specific primers can be found in **Table S1**.

### Imaging and Analysis using HCS studio 2.0 Software

Images were acquired on an ArrayScan VTi HCA Reader (ThermoFisher Scientific) using 10x or 20x magnification. For analysis, the Thermo ScienticTM TargetValidationV4.1 application was used. Readout measurements included *%* transfected cell number, fluorescence intensity, nuclear size, and nuclear shape.

### Statistical analysis

Mean as well as standard deviation (in triplicate) was used for data analysis. One way ANOVA test was used for comparison of the results. For finding the differences between groups, data was analyzed by post-Hoc, Dunnett’s multiple comparisons test. The P values of ****p<.0001; ***p<.001; **p<.01; *p<.05 were considered as statistically significant. Graph pad prism software (v.7.0) was used for data analysis.

## COLCLUSIONS

In summary, here we report the high-throughput screening and development of PBAE-based, biodegradable nanoparticles as efficient vehicles for delivering pDNA to human iPSc-RPE monolayers using a combinatorial chemistry approach. By screening a total of 140 synthesized PBAEs with varying chemical structures, we identified lead PBAE structures that resulted in markedly increased pDNA delivery efficiency both *in vitro.* Our results suggest that PBAE can effectively complex pDNA into nanoparticles, and protect the pDNA from being degraded by environmental nucleases and eventually deliver effectively to RPE monolayers. Furthermore, our results support our hypothesis that PBAE mediated pDNA delivery efficiency can be modulated by tuning PBAE end group chemistry. Using human iPSc-RPE monolayers as model cell types, we identified a few PBAE polymers that allow efficient pDNA delivery at levels that are comparable or even surpassing commercial reagents like Lipofectamine 3000 and DNA-In. Unlike lipofectamine 3000 and DNA-In, which are non-degradable, the biodegradable nature of PBAE-based nanoparticles facilitates *in vitro* applications and clinical translation. Together, our results highlight the promise of PBAE-based nanoparticles as novel nonviral gene carriers for pDNA delivery into hard-to-transfect cell RPE monolayers.

## ACKNOWLEDGEMENTS

We are grateful to Baranda S. Hansen for technical assistance. The authors thank the Wilmer Microscopy and Imaging Core Facility (EY001765) at Johns Hopkins for use of their confocal microscopy.

## AUTHOR CONTRIBUTION

Overall conceptualization, B.M., D.R.W., D.J.Z and J.J.G; Methodology and Investigation, B.M., D.R.W., S.S.R, M.P.S., C.B., and Y.R.; Resource Generation (plasmid DNA designing and production), K.J.W., (stem cell differentiation and maintenance), S.S.R.; Writing—Original Draft, B.M., D.R.W., C.B., D.J.Z and J.J.G; Writing—Review & Editing, B.M., D.R.W., C.B., D.J.Z and J.J.G.; Funding Acquisition, D.J.Z., J.J.G., S.S.R., D.R.W., and Y.R.; Supervision and Project Administration, D.J.Z and J.J.G.

## FIGURE LEGENDS

**Figure S1:** Ineffective endcap monomers. Endcap structures shown were tested and confirmed to effectively react with acrylate terminated PBAE polymer 4-4-Ac but the resulting polymers were wholly ineffective for delivery of plasmid DNA to HEK293T cells. These E-monomers were excluded from large library endcapping for transfection efficacy studies in harder-to-transfect RPE monolayers.

**Figure S2: In vitro high throughput screening of PBAE nanoparticles in subconfluent D3 RPE monolayer. (A)** Representative images showing mCherry transfected RPE cells. Heat maps showing the **(B)** percentage transfected RPE cells and **(C)** percentage survival rate following the introduction of a combinations of 140 different nanoparticles to confluent RPE monolayer at day 3 post seeding. The color scale bar refers to the percentage transfection efficiency and percentage survival that was calculated based on the number of mCherry positive cells detected from total number of cell population.

Figure S3: Base polymer PBAEs were characterized via ^1^H NMR (500 Mhz) following 2x diethyl ether precipitated to verify that base polymer structures were acrylate terminated. The ratio of integrated acrylate peak area to s-monomer carbon area was used to determine molecular weight M_N_ of base polymers. Calibration and contamination peaks include CDCl_3_ 7.26; DMSO 2.62, diethyl ether ——, tetramethyl silane (TMS) 0.

**Figure S4: in vitro transfection of confluent D25 RPE monolayer with commercial transfection reagents.** Representative Cellomics micrographs showing transfected D25 RPE cells (red) and the endogenous nuclear GFP protein (green) of RPE monolayers transfected either with **(A)** DNA-In or with **(B)** lipofectamine 3000. Histogram showing relative cell count and transfection efficacy of RPE cells transfected with **(C)** DNA-In or with **(D)** lipofectamine 3000. **(E)** Histogram showing number of cells counted over the course of RPE differentiation. Transfection efficiency shown by percentage of mCherry positive cells, quantified using a specific algorithm designed for transfection assay in a High Content Analysis platform. Confocal micrograph Scale bar: 50μm.

## REFERENCES

1. Bainbridge JW, Tan MH, Ali RR. Gene therapy progress and prospects: the eye. Gene Ther. Aug 2006;13(16):1191–1197.

2. Jia F, Wilson KD, Sun N, et al. A nonviral minicircle vector for deriving human iPS cells. Nat Methods. Mar 2010; 7(3): 197–199.

3. Nauta A, Seidel C, Deveza L, et al. Adipose-derived stromal cells overexpressing vascular endothelial growth factor accelerate mouse excisional wound healing. Mol Ther. Feb 2013;21(2):445–455.

4. Baum C, Kustikova O, Modlich U, Li Z, Fehse B. Mutagenesis and oncogenesis by chromosomal insertion of gene transfer vectors. Hum Gene Ther. Mar 2006;17(3):253–263.

5. Sasaki A, Kinjo M. Monitoring intracellular degradation of exogenous DNA using diffusion properties. J Control Release. Apr 2 2010;143(1):104–111.

6. Bitner H, Mizrahi-Meissonnier L, Griefner G, Erdinest I, Sharon D, Banin E. A homozygous frameshift mutation in BEST1 causes the classical form of Best disease in an autosomal recessive mode. Invest Ophthalmol Vis Sci. Jul 18 2011;52(8):53 32–53 38.

7. den Hollander AI, Roepman R, Koenekoop RK, Cremers FP. Leber congenital amaurosis: genes, proteins and disease mechanisms. Prog Retin Eye Res. Jul 2008;27(4):391–419.

8. Liu X, Bulgakov OV, Darrow KN, et al. Usherin is required for maintenance of retinal photoreceptors and normal development of cochlear hair cells. Proc Natl Acad Sci U S A. Mar 13 2007;104(11):4413–4418.

9. Mastrobattista E, Hennink WE. Polymers for gene delivery: Charged for success. Nat Mater. Dec 15 2011;11(1):10–12.

10. Lundstrom K. Latest development in viral vectors for gene therapy. Trends Biotechnol. Mar 2003;21(3):117–122.

11. Boylan NJ, Kim AJ, Suk JS, et al. Enhancement of airway gene transfer by DNA nanoparticles using a pH-responsive block copolymer of polyethylene glycol and poly-L-lysine. Biomaterials. Mar 2012;33(7):2361–2371.

12. Cheng W, Yang C, Hedrick JL, Williams DF, Yang YY, Ashton-Rickardt PG. Delivery of a granzyme B inhibitor gene using carbamate-mannose modified PEI protects against cytotoxic lymphocyte killing. Biomaterials. May 2013;34(14):3697–3705.

13. de la Fuente M, Ravina M, Paolicelli P, Sanchez A, Seijo B, Alonso MJ. Chitosan-based nanostructures: a delivery platform for ocular therapeutics. Adv Drug Deliv Rev. Jan 31 2010;62(1):100–117.

14. Kim TH, Park IK, Nah JW, Choi YJ, Cho CS. Galactosylated chitosan/DNA nanoparticles prepared using water-soluble chitosan as a gene carrier. Biomaterials. Aug 2004;25(17):3783–3792.

15. Read ML, Singh S, Ahmed Z, et al. A versatile reducible polycation-based system for efficient delivery of a broad range of nucleic acids. Nucleic Acids Res. May 24 2005;33(9):e86.

16. Wang H, Shi HB, Yin SK. Polyamidoamine dendrimers as gene delivery carriers in the inner ear: How to improve transfection efficiency. Exp Ther Med. Sep 2011;2(5):777–781.

17. Yu H, Russ V, Wagner E. Influence of the molecular weight of bioreducible oligoethylenimine conjugates on the polyplex transfection properties. AAPSJ. Sep 2009;ll(3):445–455.

18. Pack DW, Hoffman AS, Pun S, Stayton PS. Design and development of polymers for gene delivery. Nat Rev Drug Discov. Jul 2005;4(7):581–593.

19. Yang F, Green JJ, Dinio T, et al. Gene delivery to human adult and embryonic cell-derived stem cells using biodegradable nanoparticulate polymeric vectors. Gene Ther. Apr 2009;16(4):533–546.

20. Sunshine JC, Sunshine SB, Bhutto I, Handa JT, Green JJ. Poly(beta-amino ester)-nanoparticle mediated transfection of retinal pigment epithelial cells in vitro and in vivo. PLoS One. 2012;7(5):e37543.

21. Shmueli RB, Sunshine JC, Xu Z, Duh EJ, Green JJ. Gene delivery nanoparticles specific for human microvasculature and macrovasculature. Nanomedicine. Oct 2012;8(7):1200–1207.

22. Sunshine J, Green JJ, Mahon KP, et al. Small-Molecule End-Groups of Linear Polymer Determine Cell-type Gene-Delivery Efficacy. Adv Mater. Dec 28 2009;21(48):4947–4951.

23. Strauss O. The retinal pigment epithelium in visual function. Physiol Rev. Jul 2005;85(3):845–881.

24. Kawa MP, Machalinska A, Roginska D, Machalinski B. Complement system in pathogenesis of AMD: dual player in degeneration and protection of retinal tissue. J Immunol Res. 2014;2014:483960.

25. Wang S, Koster KM, He Y, Zhou Q. miRNAs as potential therapeutic targets for age-related macular degeneration. Future Med Chem. Mar 2012;4(3):277–287.

26. Abul-Hassan K, Walmsley R, Boulton M. Optimization of non-viral gene transfer to human primary retinal pigment epithelial cells. Curr Eye Res. May 2000;20(5):361–366.

27. Bejjani RA, BenEzra D, Cohen H, et al. Nanoparticles for gene delivery to retinal pigment epithelial cells. Mol Vis. Feb 17 2005;11:124–132.

28. Chaum E, Hatton MP, Stein G. Polyplex-mediated gene transfer into human retinal pigment epithelial cells in vitro. J Cell Biochem. Nov 1999;76(1):153–160.

29. Jayaraman MS, Bharali DJ, Sudha T, Mousa SA. Nano chitosan peptide as a potential therapeutic carrier for retinal delivery to treat age-related macular degeneration. Mol Vis. 2012;18:2300–2308.

30. Liu HA, Liu YL, Ma ZZ, Wang JC, Zhang Q. A lipid nanoparticle system improves siRNA efficacy in RPE cells and a laser-induced murine CNV model. Invest Ophthalmol Vis Sci. Jul 1 2011;52(7):4789–4794.

31. Mannermaa E, Ronkko S, Ruponen M, Reinisalo M, Urtti A. Long-lasting secretion of transgene product from differentiated and filter-grown retinal pigment epithelial cells after nonviral gene transfer. Curr Eye Res. May 2005;30(5):345–353.

32. Mannisto M, Ronkko S, Matto M, et al. The role of cell cycle on polyplex- mediated gene transfer into a retinal pigment epithelial cell line. J Gene Med. Apr 2005;7(4):466–476.

33. Mannisto M, Vanderkerken S, Toncheva V, et al. Structure-activity relationships of poly(L-lysines): effects of pegylation and molecular shape on physicochemical and biological properties in gene delivery. J Control Release. Sep 18 2002;83(1):169–182.

34. Peeters L, Sanders NN, Jones A, Demeester J, De Smedt SC. Post-pegylated lipoplexes are promising vehicles for gene delivery in RPE cells. J Control Release. Aug 28 2007;121(3):208–217.

35. Peng CH, Cherng JY, Chiou GY, et al. Delivery of Oct4 and SirT1 with cationic polyurethanes-short branch PEI to aged retinal pigment epithelium. Biomaterials. Dec 2011;32(34):9077–9088.

36. Gan Q, Wang T, Cochrane C, McCarron P. Modulation of surface charge, particle size and morphological properties of chitosan-TPP nanoparticles intended for gene delivery. Colloids Surf B Biointerfaces. Aug 2005;44(2- 3):65–73.

37. Vercauteren D, Piest M, van der Aa LJ, et al. Flotillin-dependent endocytosis and a phagocytosis-like mechanism for cellular internalization of disulfide-based polyfamido amine)/DNA polyplexes. Biomaterials. Apr 2011;32(ll):3072–3084.

38. Klimanskaya I, Hipp J, Rezai KA, West M, Atala A, Lanza R. Derivation and comparative assessment of retinal pigment epithelium from human embryonic stem cells using transcriptomics. Cloning Stem Cells. 2004;6(3):217–245.

39. Molla MR, Levkin PA. Combinatorial Approach to Nanoarchitectonics for Nonviral Delivery of Nucleic Acids. Adv Mater. Feb 10 2016;28(6):1159–1175.

40. Frohlich E. The role of surface charge in cellular uptake and cytotoxicity of medical nanoparticles. Int J Nanomedicine. 2012;7:5577–5591.

41. Tomita Y, Rikimaru-Kaneko A, Hashiguchi K, Shirotake S. Effect of anionic and cationic n-butylcyanoacrylate nanoparticles on NO and cytokine production in Raw264.7 cells. Immunopharmacol Immunotoxicol. Dec 2011;33(4):730–737.

42. Bishop CJ, Ketola TM, Tzeng SY, et al. The effect and role of carbon atoms in poly(beta-amino ester)s for DNA binding and gene delivery. J Am Chem Soc. May 8 2013;135(18):6951–6957.

43. Tzeng SY, Wilson DR, Hansen SK, Quinones-Hinojosa A, Green JJ. Polymeric nanoparticle-based delivery of TRAIL DNA for cancer-specific killing. Bioeng Transl Med. Jun 2016;1(2):149–159.

44. Maruotti J, Wahlin K, Gorrell D, Bhutto I, Lutty G, Zack DJ. A simple and scalable process for the differentiation of retinal pigment epithelium from human pluripotent stem cells. Stem Cells Transl Med. May 2013;2(5):341–354.

45. Maruotti J, Sripathi SR, Bharti K, et al. Small-molecule-directed, efficient generation of retinal pigment epithelium from human pluripotent stem cells. Proc Natl Acad Sci U S A. Sep 1 2015;112(35):10950–10955.

46. Gamm DM, Melvan JN, Shearer RL, et al. A novel serum-free method for culturing human prenatal retinal pigment epithelial cells. Invest Ophthalmol Vis Sci. Feb 2008;49(2):788–799.

